# Systematic analysis of intrinsic enhancer-promoter compatibility in the mouse genome

**DOI:** 10.1101/2021.10.21.465269

**Authors:** Miguel Martinez-Ara, Federico Comoglio, Joris van Arensbergen, Bas van Steensel

## Abstract

Gene expression is in part controlled by cis-regulatory elements (CREs) such as enhancers and repressive elements. Anecdotal evidence has indicated that a CRE and a promoter need to be biochemically compatible for promoter regulation to occur, but this compatibility has remained poorly characterised in mammalian cells. We used high-throughput combinatorial reporter assays to test thousands of CRE – promoter pairs from three Mb-sized genomic regions in mouse cells. This revealed that CREs vary substantially in their promoter compatibility, ranging from striking specificity for a single promoter to quantitative differences in activation across a broad set of promoters. More than half of the tested CREs exhibit significant promoter selectivity. Housekeeping promoters tend to have similar CRE preferences, but other promoters exhibit a wide diversity of compatibilities. Higher-order TF motif combinations may account for compatibility. CRE–promoter selectivity does not correlate with looping interactions in the native genomic context, suggesting that chromatin folding and compatibility are two orthogonal mechanisms that confer specificity to gene regulation.

## INTRODUCTION

How genes are regulated by cis-regulatory elements (CREs) such as enhancers and repressor elements is a long-standing topic in molecular biology [1–10]. One conundrum is how CREs ‘choose’ their target promoters. Some enhancers can activate multiple promoters *in cis* over short and long genomic distances [11–13], while others show remarkable specificity, regulating only one of its neighbouring promoters or even skipping one or more promoters to activate more distal ones. In part, 3D folding and compartmentalisation of the chromatin fibre help to establish this specificity, by facilitating certain enhancer-promoter contacts and curbing others [12–14].

However, there is also substantial evidence that biochemical (in)compatibility between CREs and promoters contributes to the specificity of their regulatory interactions. This is akin to a lock-and-key mechanism: proteins bound to the CRE and the promoter must be compatible in order to form a productive complex. Examples of such intrinsic selectivity have been documented particularly in *Drosophila*, and in some instances could be attributed to a specific sequence motif in the promoter [15–19]. Data obtained with massively parallel reporter assays (MPRAs) in *Drosophila* cells have suggested a general separation of enhancer-promoter compatibility into housekeeping and tissue-specific classes [20]. Some of this specificity may be determined by the recruitment of co-factors [21]. However, a thorough understanding of the underlying mechanisms is still lacking.

While several studies of individual enhancer-promoter combinations indicate that biochemical compatibility also plays a role in mammals (e.g., [22–26]), systematic studies of this mechanism have so far been lacking in mouse or human cells. Thus, it is still unknown how widespread such intrinsic compatibility is in mammalian cells, and what drives this compatibility.

In order to address this issue, we systematically tested the compatibility of thousands of combinations of candidate CREs (cCREs) and promoters using MPRAs. We used plasmid-based MPRAs because they are highly scalable [27–29], and because episomal plasmids provide an isolated context that minimises confounding effects of variable chromatin environments and differences in 3D folding. However, so far MPRAs have mostly been used to assess the activity of single elements, either as enhancers or as promoters [27, 30–34], except for one recent study that tested combinations of synthetic elements [29]. To be able to dissect compatibility between enhancers and promoters systematically, we designed cloning strategies that allowed us to test thousands of pairwise cCRE–promoter combinations in different positions and orientations in a reporter plasmid.

As models, we chose three genomic loci of 1-3 Mb in mouse embryonic stem cells (mESCs). From these loci, which each encompass ~20 genes, we tested a large fraction of all possible pairwise cCRE–promoter (cCRE-P) combinations. We found that more than half of the active cCREs exhibit significant selectivity for specific subsets of promoters. We dissected some of the underlying sequence determinants. Furthermore, we provide evidence suggesting that 3D folding and intrinsic compatibility are independent mechanisms. Our experimental strategy and datasets provide novel insights into the logic and mechanisms of cCRE-promoter specificity.

## RESULTS

### Experimental design

To maximise the probability of testing biologically relevant enhancer-promoter pairs, we combined cCREs and promoters coming from the same region in the genome. We selected three loci of 1-3 Mb in size, each roughly centred around a gene (*Nanog*, *Tfcp2l1* or *Klf2*) that is key to the control of pluripotency of mESCs. The regulation of these genes is still incompletely understood. In addition, each locus contains about 20 other genes (**Figure 1A-C**).

**Figure 1.**
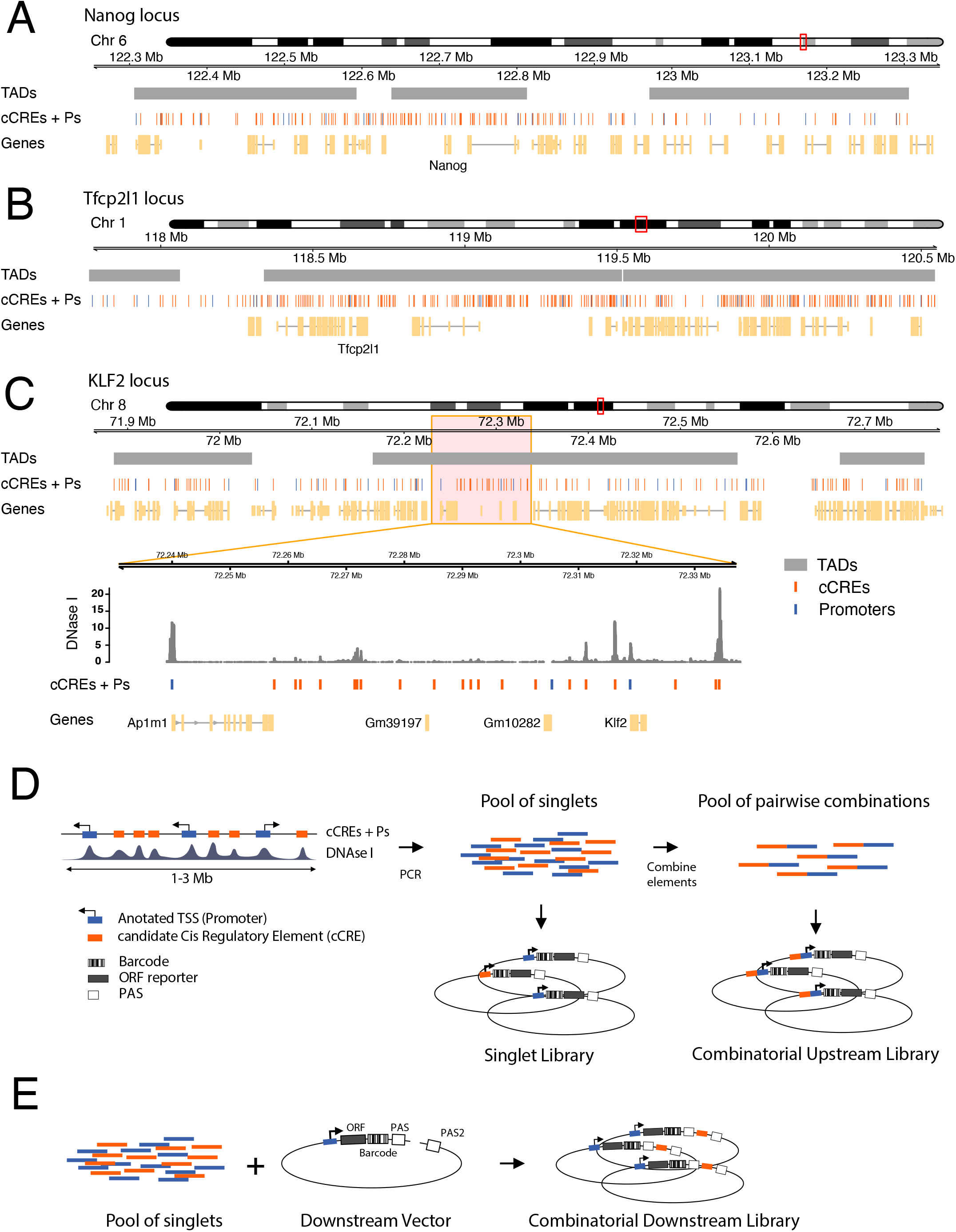
Regulatory element selection and library construction. **A-C)** Representations of *Nanog*, *Tfcp2l1*, and *Klf2* loci, respectively. In **C)** the zoom-in displays a DNAse I sensitivity track [38] where peaks overlap with cCREs. **D)** Cloning strategy for the Upstream assay. cCREs and promoters were amplified by PCR from genomic DNA and pooled. Fragments in this pool were then randomly ligated to generate duplets. Singlets and duplets were cloned into the same barcoded vector to generate two libraries per locus, a singlet library and a combinatorial library. **E)** Cloning strategy for the Downstream assay. The singlet pool from the *Klf2* locus was cloned into ten vectors, each of them carrying a different promoter. The resulting ten sub-libraries were combined into one Downstream assay library.

For promoters in the regions of interest we included approximately the −350 to +50 bp segments around all GENCODE-annotated [35] transcription start sites (TSSs). The choice to focus on the range −350 to +50 bp was motivated by our previous study of human promoters, which indicated that most of the relevant information for promoter function is generally contained within this range [30]. This definition of promoters is longer than that of core promoters (which are usually only ~100 bp long) as was used in most previous enhancer reporter assays [21, 27, 29, 32–34, 36]. We considered this to be important, because the extra regulatory information contained in those additional sequences may be relevant for interactions of the promoters with CREs.

Compared to promoters, the annotation of cCREs is much less accurate. However, most cCREs are centred around DNase I hypersensitive sites (DHS) [5, 37, 38]. We therefore selected fragments of ~400 bp centred around all detected DHS peaks in each locus (**Figure 1A-C**). This definition of cCREs within the range of typical enhancer definitions [39]. Some authors consider enhancers combinations of multiple DHSs or longer stretches of DNA sequences. However, other studies have shown that the activity of these long enhancers can be reproduced by shorter versions of ~500 bp [40, 41]. Coordinates of all tested genomic fragments are provided in **Supplementary Dataset 1**.

We designed two MPRA variants to test many cCRE-P combinations (**Figure 1D-E**). In the first variant, which we will refer to as Upstream assay, we obtained 82-192 individual cCREs and 18-25 P elements per locus by PCR amplification (**Table 1)**. We pooled all of these fragments and randomly ligated them to form dimer fragments, which we then cloned *en masse* into a reporter vector, *upstream* of a randomly barcoded transcription unit that lacked a promoter itself. This resulted into highly complex libraries of cCRE-P, cCRE-cCRE, P-P and P-cCRE pairs, with each individual element in two possible orientations. We then sequenced the libraries to identify the paired fragments, their orientations in the reporter vector, and their linked barcodes. Owing to the simple random ligation step, libraries with tens of thousands of cCRE-P combinations can be obtained with this approach (**Table 1** and **Supplementary Table 1)**. Here, we focus on the analysis of cCRE-P pairs, but data from all other configurations are also provided as **Supplementary Dataset 2**.

**Table 1.** Numbers of tested Promoters (Ps), cCREs and cCRE–P pairs in each combinatorial MPRA library.

| Library | Ps present | cCREs present | cCRE–P pairs tested | cCRE–P pairs (orientation-independent) |
| --- | --- | --- | --- | --- |
| Klf2 Upstream | 23 | 82 | 3758 | 1400 |
| Nanog Upstream | 18 | 88 | 1321 | 595 |
| Tfcp2l1 Upstream | 25 | 198 | 5599 | 2490 |
| Klf2 Downstream | 10 | 84 | 1364 | 752 |

In a second and complementary approach, we constructed a library in which the cCREs are placed *downstream* of the reporter gene, i.e., separated ~1kb from the promoter (**Figure 1E**). This was done in two steps: we first cloned a selection of 10 promoters upstream of the barcoded transcription unit, resulting in a set of reporters with different promoters. Next, we inserted a pool of cCREs into this set, downstream of the barcoded reporter unit and in both possible orientations. We will refer to the assays done with the resulting library as Downstream assay. Due to the two-step cloning protocol, the Downstream assay is less scalable than the Upstream assay, but nevertheless allows for testing of hundreds of cCRE–P combinations (**Table 1**).

We used all P and cCRE DNA fragments from each of the three loci in separate Upstream assays, whereas we focused on ten promoters and all cCREs from the *Klf2* locus in the Downstream assay. **Table 1** provides summary statistics of the individual library compositions. Due to the random nature of the combinatorial cloning, we did not recover all possible pairs. Nevertheless, in the three Upstream assays combined we tested a total of 10,678 cCRE-P pairs, or 3,747 pairs if we do not take orientations into account. For the Downstream assay these numbers were 1,364 and 752, respectively. From the *Klf2* locus 847 and 676 pairs, respectively, overlapped between the Upstream and Downstream assay. As references, we also inserted each P and cCRE individually (i.e., unpaired) in the upstream position.

### Boost indices estimate promoter-specific activity of cCREs

We then transiently transfected each of these libraries into mESCs. Twenty-four hours after transfection we collected mRNA from the cells, and counted the transcribed barcodes by reverse transcription followed by PCR amplification and high-throughput sequencing. In parallel, barcodes were counted in the plasmid libraries. For each barcode we then normalised the counts in cDNA over the counts detected in the plasmid DNA. Further data processing is described in the Methods. We performed 3 biological replicates per library, which correlated with an average Pearson r=0.87 (0.83 to 0.90) for the Upstream assay and r=0.98 (0.98 to 0.99) for the Downstream assay. (**Figure S1 A-C**)

We first analysed the transcriptional activities of all singlet (unpaired) P and cCREs in the upstream position. For promoters, these basal activities varied over a ~100-fold dynamic range (**Figure 2A**; **Figure S2A**). Of all cCREs, 40.4% showed detectable transcriptional activity in the upstream position without any P (**Figure 2A**; **Figure S2A**). Such autonomous transcriptional activity is a frequently observed property of enhancers [30, 42, 43], and hence these elements are likely to be enhancers. For a few cCREs this activity was as high as some of the strongest promoters, suggesting that they may in fact be un-annotated promoters or very strong enhancers.

**Figure 2.**
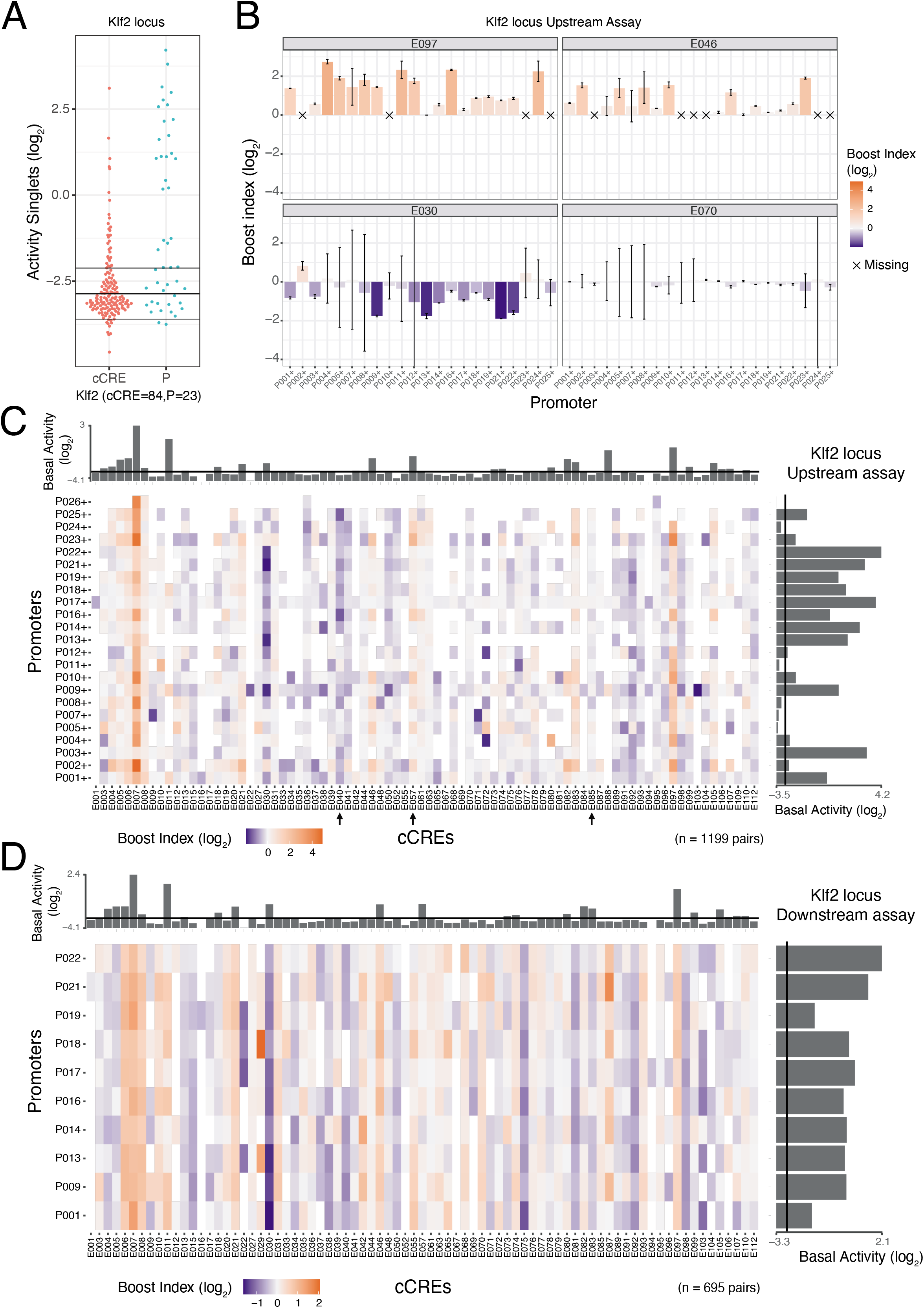
Singlet and combinatorial activities of cCREs and promoters from the Klf2 locus. **A)** Transcription activities of singlet cCREs and promoters. Each dot represents the mean activity of one singlet. Horizontal lines represent the average background activity of empty vectors (black line) plus or minus two standard deviations (grey lines). Elements with activities more than two standard deviations above the average background signal are defined as active. **B)** Examples of Upstream assay cCRE-P combinations for cCREs E097, E046, E030 and E070 of the *Klf2* locus. Barplots represent the mean boost index of each combination, vertical lines represent the standard deviations. Crosses mark missing data. **C-D)** Boost index matrices of cCRE–P combinations from the *Klf2* locus according to Upstream **(C**) and Downstream **(D)** assays. White tiles indicate missing data. Barplots on the right and top of each panel show basal activities of each tested P or cCRE, respectively, with the black line indicating the background activity of the empty vector. All data are averages over 3 independent biological replicates.

We then determined the ability of each cCRE to alter the activity of each linked P. For this, we calculated a *boost index* for each cCRE–P pair, defined as the log_2_-fold change in activity of the cCRE-P pair compared to the P element alone. Unexpectedly, 20 negative controls that we included in the *Klf2* libraries, consisting of randomly generated DNA sequences of similar size and G/C content as the cCREs, showed a modestly negative boost index (median value −0.45 when inserted upstream) (**Figure S1D**). This is possibly because lengthening of the reporter constructs alters the topology, supercoiling, transfection efficiency or a combination of these parameters. We therefore corrected all cCRE–P boost indices for this non-specific negative bias (see Methods). After this correction the negative controls had a marginal residual bias (median log_2_ value −0.19), which we deemed acceptable (**Supplementary Figure S1D**).

### Identification of activating and repressive cCREs

For each of the three genomic loci, the matrix of corrected boost indices shows a wide diversity of patterns across the cCREs. We observed this both in the Upstream and Downstream assays (**Figure 2B-D**, **Supplementary Figure S2B-D**). For example, in the *Klf2* locus Upstream assay, cCRE E097 activates most of the tested promoters, while E046 (**Figure 2B**) and E057 (arrow in **Figure 2C**) only activate a distinct subset of promoters. Several elements are primarily acting as repressors (e.g, E030 (**Figure 2B**) and E040, (arrow in **Figure 2C**)), and some seem neither activating nor repressive (e.g., E070 (**Figure 2B**) and E085 (arrow in **Figure 2C**)).

We broadly classified the cCREs according to their overall effects on the linked promoters (**Figure S3A**). In the Upstream assays, 21% of cCREs showed positive boost indices that were significantly higher than the rest of cCREs across all tested promoters, indicating that they can act as enhancer elements. About 17% of the cCREs showed negative boost indices significantly below the rest of cCREs, and hence are putative repressor elements. For the remaining 62% of cCREs the boost indices across their linked promoters were not significantly higher or lower than the rest; these “ambiguous” elements either have no regulatory effects at all, or they have a mixed repressive/activating/inactive effect that depends on the linked P (see below).

We were somewhat surprised to identify similar numbers of putative enhancers and repressors, because most annotated cCREs in mammalian genomes are predicted to be enhancers rather than repressive elements [5, 44]. In some cases this repression may be underestimated in our analysis, as the estimates of negative boost indices for lowly active promoters are less reliable due to the higher noise-to-mean ratios at low expression levels (**Figure S3B**).

For activating elements, the boost indices varied in part according to the basal activities of the cCRE and promoters. Strong boosting occurred primarily at promoters with low basal activities, while highly active promoters were more difficult to boost (**FigS3C**). This suggests a saturation effect, or it could indicate that promoters with high basal activity are less dependent on distal enhancers. For cCREs, their basal activity is generally a strong positive predictor of their enhancer potency (**Fig S3D**). However, exceptions to this rule occur, as some cCRE-P pairs show high boost indices even though the basal activity of the cCRE is low (**Fig S3D**, upper left quadrant).

### cCRE effects are predominantly orientation- and position-independent

Next, we asked whether the ability of cCREs to regulate the linked promoters was generally independent of their orientation and position. This was originally posited for enhancers [1], and in some cases also reported for repressive elements [10]. Indeed, in the Upstream assays we found a general positive correlation of the boost indices between the two orientations of the cCREs (Pearson’s r=0.68) (**Figure S4A**). These results are similar to those recently obtained with a minimal core promoter [32]. In the Downstream assay the correlation between orientations was somewhat lower (Pearson’s r=0.47) (**Figure S4B**). This may be due to the lower dynamic range of the Downstream assay data (**Figure S1C**). To simplify, for all other analyses we combined the boost indices of + and - orientations of the cCREs by averaging.

We then investigated the degree of position-independence, by comparing the overlapping P-cCRE pairs from the *Klf2* locus Downstream and Upstream assays. This showed an overall Pearson correlation of 0.64 (**Figure S4C**). We conclude that repressive and activating effects of cCREs are substantially but not completely position-independent, at least for the ten tested promoters from the *Klf2* locus.

### Extensive selectivity of cCREs for promoters

Visual inspection of the boost index matrices suggested that some cCREs alter the expression of most promoters to similar degrees, while others selectively alter the expression of a subset of promoters. In addition to the examples in **Figure 2B** from the *Klf2* locus, strikingly specific promoter responses to some cCREs are illustrated for the *Tfcp2l1* locus in **Figure 3A**. For example, E060, which forms part of an annotated super-enhancer [45], activates most of the tested promoters, but with boost indices that can vary >50-fold between promoters. Two other remarkable examples from the *Tfcp2l1* locus are E091 and E096, which each activate only a single, distinct promoters out of the 11-12 promoters that were tested in each instance. Much broader specificity is observed for E064, E073, E074 and E090 from the *Nanog* locus, which are part of previously identified super-enhancers [46] (**Figure S2D**).

**Figure 3.**
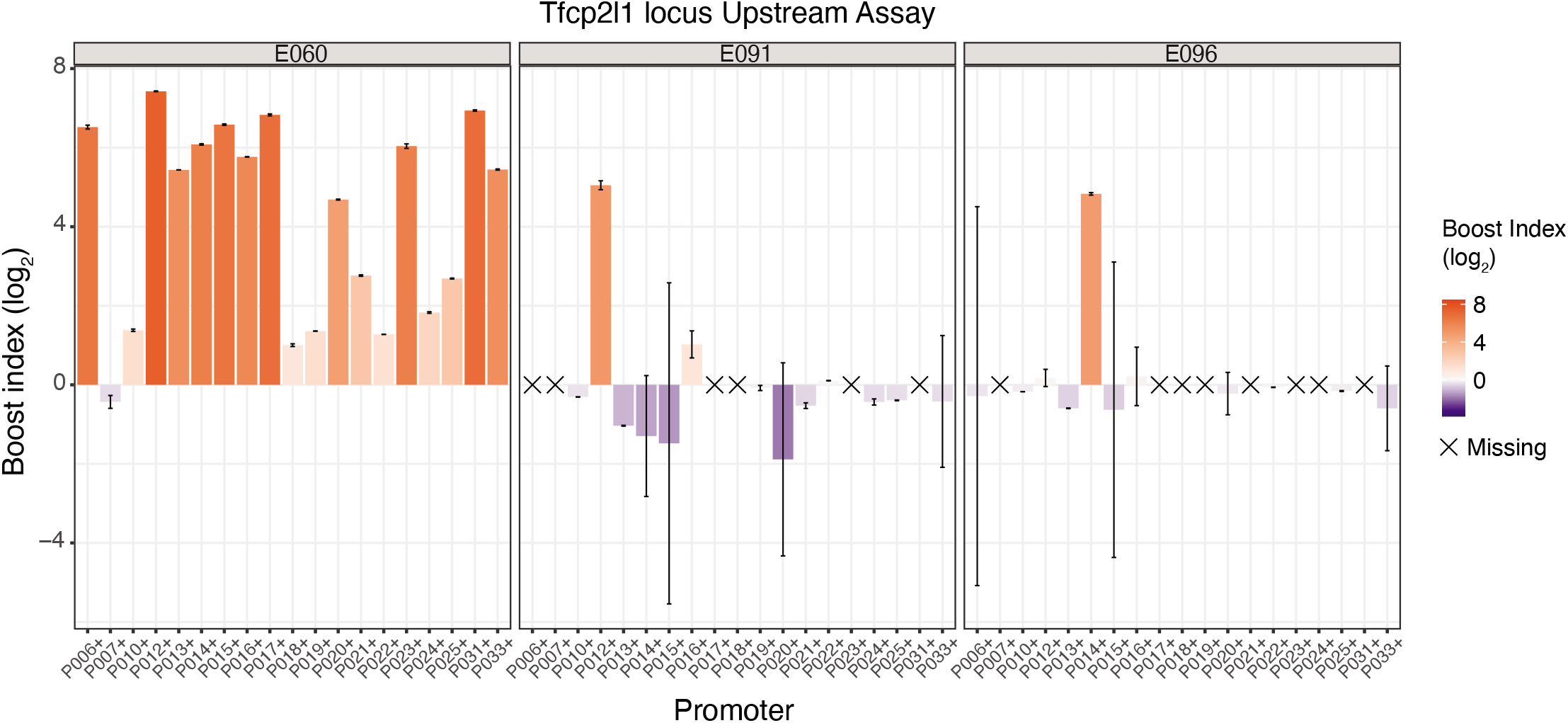
Examples of selective cCREs from the *Tfcp2l1* locus. Boost indices obtained in the Upstream assay are shown for cCRE-P combinations of cCREs E060, E091 and E096 of the *Tfcp2l1* locus. Barplots indicate the mean boost index of each combination, vertical lines indicate standard deviations. All data are averages over 3 independent biological replicates.

We investigated the degrees of selectivity more systematically. **Figure 4A-B** depicts the distribution of the boost indices for each cCRE. Clearly, some cCREs have a much broader range of boost indices than others. We used an ANOVA approach with Welch F-test to systematically identify cCREs for which the variance of boost indices was larger than could be explained by experimental noise (see methods). Strikingly, out of 233 cCREs with more than 5 tested cCRE-P combinations, a total of 139 (59.9%) (**Figure 4B-C**) showed significant unexplained variance at an estimated false-discovery rate (FDR) cutoff of 5%. Thus, at least throughout the three loci that we tested, cCRE-P selectivity is widespread, ranging from strong specificity for one or a few promoters to low specificity as seen in quantitative differences in the regulation of a broad set of promoters.

**Figure 4.**
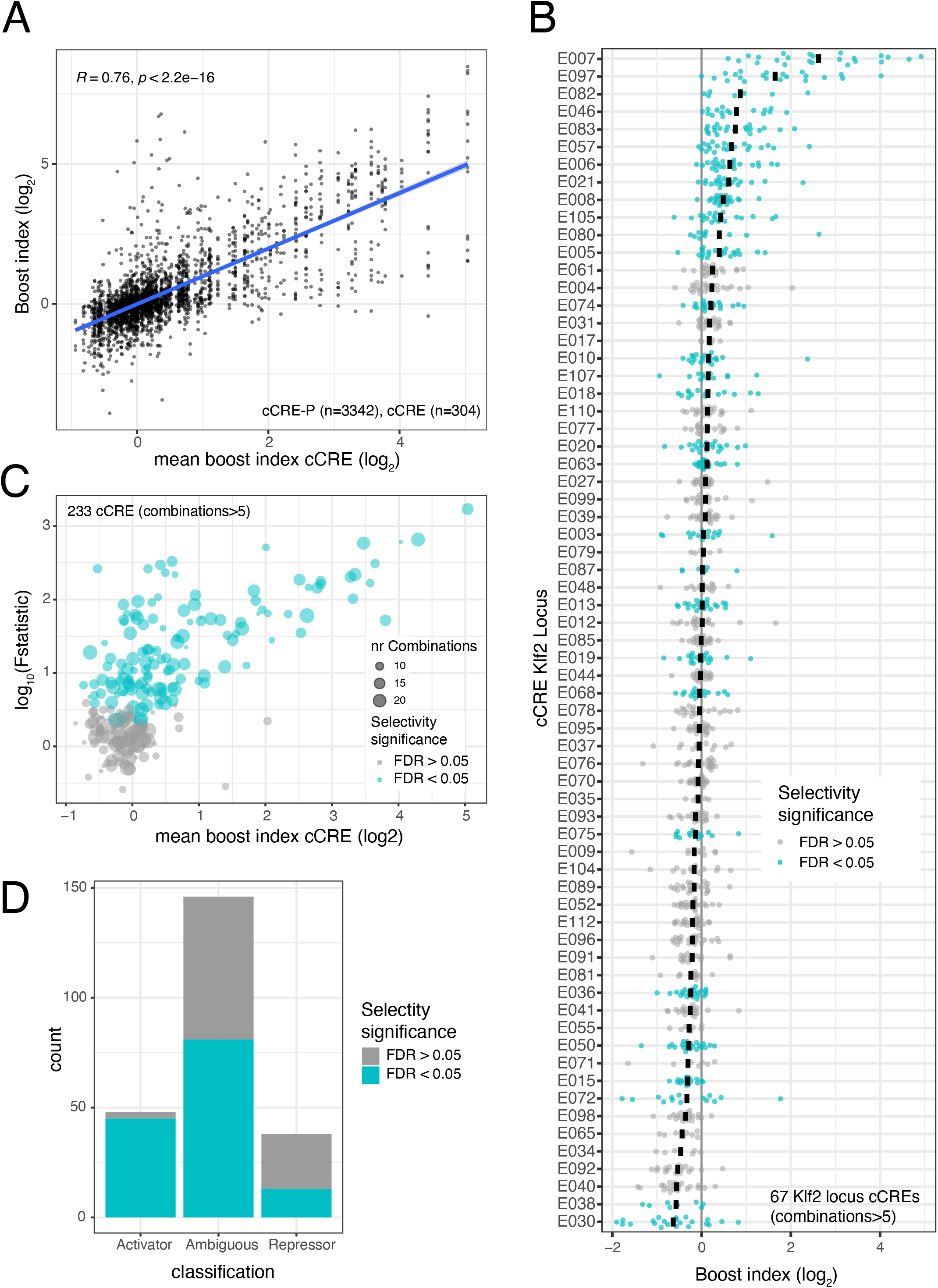
Promoter selectivity of cCREs. **A)** Plot showing the broad diversity of boost indices of many cCREs. Data are from Upstream assays of *Klf2, Nanog* and *Tfcp2l1* loci combined. Vertical axis indicates boost indices of all tested cCRE–P pairs, which are horizontally ordered by the mean boost index of each cCRE. **B)** Boost index distributions for each cCRE from the *Klf2* locus (Upstream assay). Each dot represents one cCRE–P combination; black bar represents the mean. Turquoise colouring marks cCREs that have a larger variance of their boost indices than may be expected based on experimental noise, according to the Welch F-test after multiple hypothesis correction (5% FDR cutoff). **C)** Summary of Welch F-test selectivity analysis results for all cCREs from the three loci with more than 5 cCRE–P combinations. Each dot represents one cCRE; the size of the dots indicates the number of cCRE–P pairs. Significantly selective cCREs (5% FDR cutoff) are highlighted in turquoise. **D)** Proportion of significantly selective (turquoise) cCRE in the three categories as shown in **Figure S3A**. All data are averages over 3 independent biological replicates.

Intersection of the ANOVA-based classification of selective/unselective cCREs with the above broad classification into enhancers and repressors indicates that almost all (94%) general enhancer elements exhibit significant P selectivity. In contrast, only 34% of the repressors are detectably biased towards a subset of promoters (**Figure 4D**). However, we note that this percentage may be underestimated, because at low expression levels the noise levels are higher (**Figure S3B**). Interestingly, among the “ambiguous” cCREs, 55% are in fact selective. Such elements mostly activate or repress only very few promoters (e.g., E091 and E096 from the *Tfcp2l1* locus; **Figure 3**) and leave all other promoters unaffected. The remainder of the ambiguous cCREs are probably not functional (e.g., E70 from the *Klf2* locus, **Figure 2B**). In summary, these results indicate that more than half of all tested cCREs exhibits significant preference for specific promoters.

Promoters of housekeeping and developmental genes in *Drosophila* were reported to have distinct specificities toward cCREs [47]. To investigate whether such a dichotomy could also be observed in our data, we focused on the *Klf2* locus, which has roughly equal numbers of housekeeping and non-housekeeping promoters [48] (the *Tfcp2l1* and *Nanog* loci have only three and zero housekeeping genes, respectively). Indeed, hierarchical clustering of the boost index matrix showed a rough separation of the two classes of promoters (**Figure S5A**). However, this is largely due to the highly similar cCRE specificities among the housekeeping promoters, whereas the cCRE specificities of the non-housekeeping promoters are much more diverse and generally as distinct from each other as from the housekeeping promoters (**Figure S5B**). To test whether a housekeeping versus non-housekeeping dichotomy may largely explain our identification of cCREs with significant selectivity (**Figure 4B-C**), we repeated this analysis after removing all housekeeping promoters. This yielded highly similar results (123 of 221 cCREs are significantly selective at 5% FDR cutoff, **Figure S5C**). We conclude that housekeeping promoters may be similarly regulated, but cCRE selectivity goes beyond a simple distinction between housekeeping and non-housekeeping promoters.

### Selectivity may be mediated by combinations of multiple TF motifs

Taken together, these results point to a broad spectrum of cCRE specificities for promoters, ranging from largely indiscriminate to highly selective. We searched for sequence motifs that may account for these effects, focusing on binding motifs of transcription factors (TFs) that are expressed in mESCs.

We first searched for TF motifs in the cCREs that correlate with boost indices across all promoters. This yielded several dozens of TFs that are candidate activators or repressors (**Figure S6A**). Several of these, such as Sox2, Nanog, ETV4 and GABPA are known key regulators in mESC cells [49–51]. These TFs may broadly contribute to enhancer activity.

Next, we searched for motifs associated with cCRE–P selectivity. We reasoned that selectivity may be due to certain combinations of TFs bound to cCRE and P. First, we asked whether for any TF the simultaneous presence of its motif at cCRE and P correlated with boost indices (**Figure 6SB**). This only yielded a weak association of FOXO motifs (at a 5% FDR cutoff). Possibly this is due to FOXO1, a known regulator in mESCs [52]. We then asked if selectivity may be mediated by multiple TFs rather than single TFs. For this purpose, we took the TF motifs associated with enhancer activity with effect sizes >0.1 (n=66) and searched for combinations of motifs that would be associated with higher boost indices if present at both the cCRE and the P (**Figure 5A-B**). This yielded a few dozen stronger associations (at a 1% FDR cutoff). Some of these associations may be redundant either because of motif similarity or because of motif co-occurrence. For example, the 5 associations between Sox2 and Klf motifs may represent the Klf4-Sox2 pair (**Figure 5B**) which are known to cooperate in mESCs [53]. These results indicate that selectivity may be mediated by combinations of multiple TF motifs. Our dataset does not provide sufficient statistical power for an exhaustive search of such combinations.

**Figure 5.**
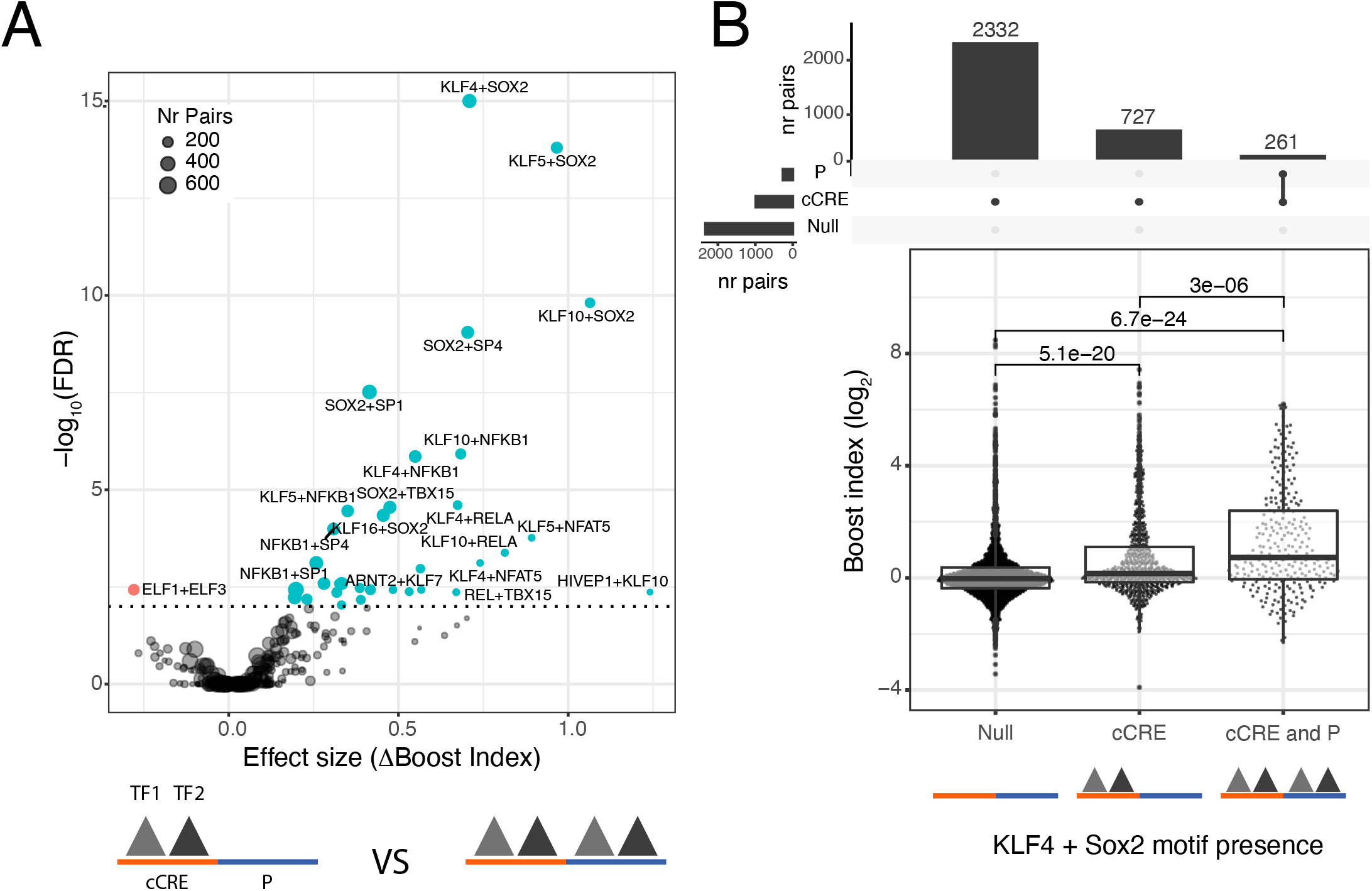
Association of TF motif Duos with higher boost indices. **A)** Results of TF survey for self-compatible TF motif Duos. TF motif duos associated with higher or lower boost indices at a 1% FDR cutoff are highlighted. **B)** Association of Sox2+Klf4 motifs at both cCRE and P with higher boost indices. cCRE-P combinations are split into 3 groups according to presence or absence of Sox2+Klf4 motifs both at the cCRE and the promoter, or only the cCRE. Numbers at the top of horizontal brackets are the p-values obtained from comparing the different groups boost index distributions using a Wilcoxon rank-sum test. Boxplots represent median and interquartile ranges. Barplots at the top represent the number of combinations in each group.

### Chromatin looping is independent of compatibility

Finally, we considered that certain pairs of cCREs and promoters frequently contact each other in the nucleus, as is indicated by focal or stripe-like enrichment patterns in high-resolution Hi-C maps [54, 55]. While long-range contacts are irrelevant in our MPRAs because the tested elements are directly linked, we asked whether such physical contacts in the native genomic context are related to the selectivity of cCREs for certain promoters according to our MPRAs. We considered two models. In one model, the biochemical interactions that underlie cCRE-P selectivity may promote or stabilise cCRE-P looping interactions. Alternatively, looping interactions and cCRE-P selectivity may be independent aspects of cCRE-P interplay that each work by different mechanisms.

To discriminate between these two models, we investigated whether the boost indices of cCRE-P pairs correlate with their contact frequencies in Micro-C, a high-resolution variant of Hi-C [55]. Remarkably, we found no correlation between these two quantities (**Figure 6A**). We also found an extremely weak, although statistically significant, correlation between higher boost indices and longer linear distances of cCRE-P pairs along the genome (**Figure 6B**).

**Figure 6.**
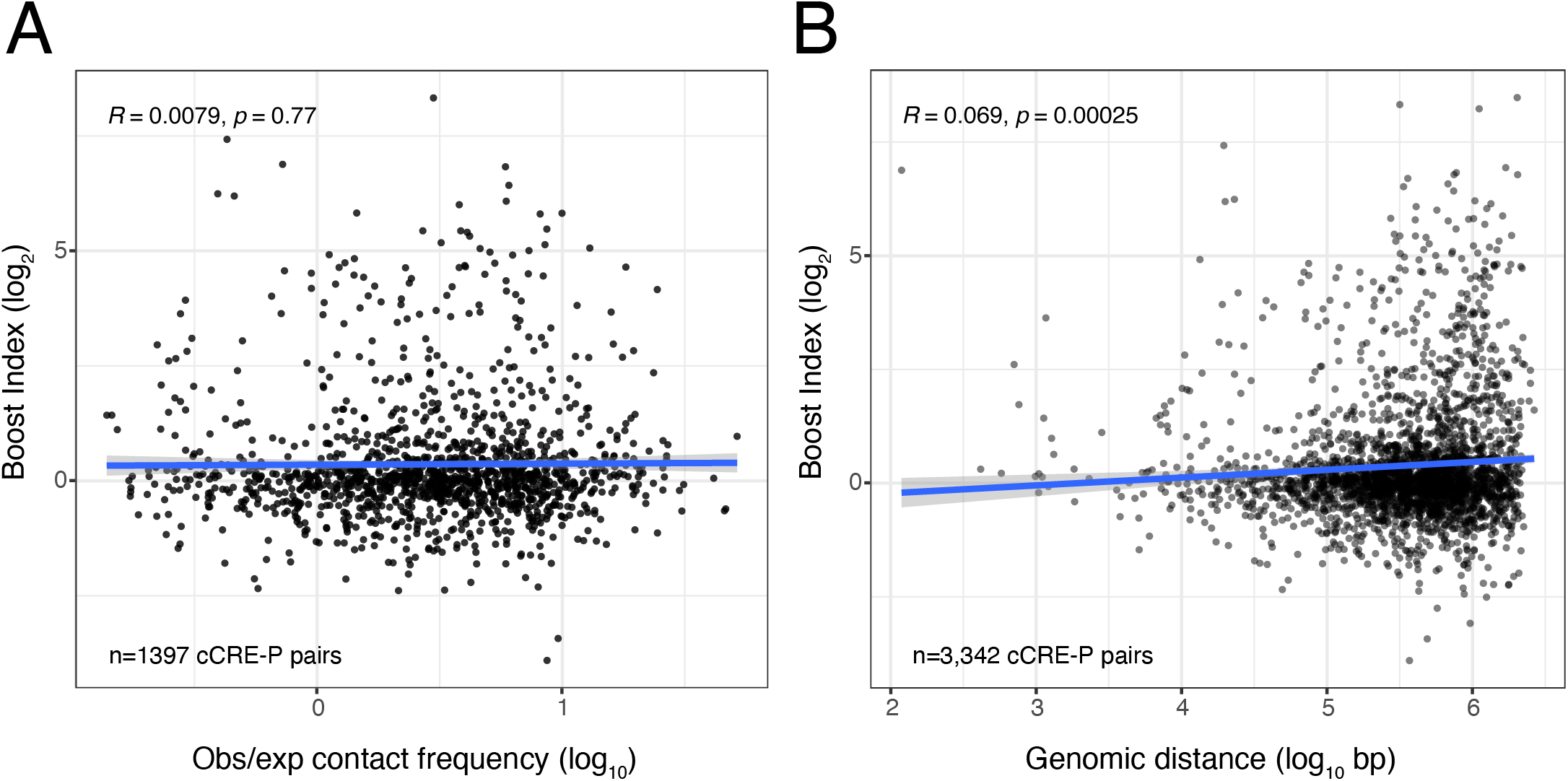
Absent or very weak correlation between boost indices and **(A)** contact frequencies according to micro-C [55] or **(B)** linear genomic distance, for all cCRE-P pairs from the three loci combined. All boost index data are averages over 3 independent biological replicates.

We conclude that cCRE-P contacts in the nucleus may be independent of their functional compatibility as detected in our reporter assays, raising the interesting possibility that chromatin looping and compatibility are two orthogonal mechanisms of gene regulation.

## DISCUSSION

Only a few other studies have so far attempted to analyse cCRE–P compatibility systematically. An early survey of 27 cCRE–P combinations in human cells did not find evidence for specificity [56], but the assay employed may have been insufficiently quantitative, and the choice of tested elements may have been biased. In contrast, testing of ~200 cCRE–P pairs in zebrafish pointed to extensive specificity [57]. An MPRA study in *Drosophila* cells using seven different promoters and genome-wide cCREs suggested that cCRE–P specificity broadly separates between housekeeping and tissue-specific promoters [47]. To our knowledge, our systematic combinatorial testing of cCRE–P combinations in mESCs is the first large-scale study in mammalian cells. The results reveal a broad spectrum of specificities: some cCREs are promiscuous, others are highly specific for certain promoters, and in many instances the specificity is quantitative rather than qualitative. By statistical analysis we found that more than half of the cCREs exhibit a degree of specificity that cannot be explained by experimental noise.

It is likely that cCRE–P compatibility is governed by a complex grammar of TF combinations. Underlying this grammar may be a diversity of molecular mechanisms, including direct and indirect TF-TF interactions [e.g., 53], local concentration of activating factors [33, 58], or functional bridging by cofactors [21, 59]. Due to the complexity of this grammar, its elucidation may require much larger cCRE–P combinatorial datasets than generated here, as well as systematic mutational analysis [60, 61] of individual cCRE–P combinations. Nevertheless, our statistical analysis highlights several candidate combinations of TF motifs that may contribute to the compatibility of some cCRE–P pairs.

Our data indicate that some of the cCREs tested may be repressive elements rather than enhancers even though they were selected from DHSs. This is similar to a recent screen of cCREs in human cells, which identified a large set of candidate repressive elements [62] and to another screen in *Drosophila* [63]. It will be interesting to further explore the physiological regulatory role of these elements. Particularly to understand their influence on close genes and how repression works in open regions of the genome.

Surprisingly, we found that the boost indices of cCRE–P pairs generally do not correlate with their contact frequencies in the native chromatin context. This suggests that 3D genome organisation and compatibility are regulated by different mechanisms. We envision that compatibility and 3D organisation may be two independent layers necessary for correct selective gene regulation: 3D organisation such as the formation of chromatin loops and compartments may determine whether CREs and promoters are able to interact, while compatibility may determine whether such an interaction is functional, i.e., gives rise to a change in P activity.

Our current data were generated with transiently transfected plasmids. Advantages of this approach are that it largely eliminates possible confounding effects of chromatin packaging and 3D folding, and that thousands of cCRE–P combinations could be tested. Even higher throughput combinatorial MPRAs will be useful in order to fully dissect the rules behind compatibility either by testing more cCRE-P combinations or mutagenised cCRE-P pairs. However, further studies are needed to verify and analyse the impact of the observed specificities in the native genomic context. Due to genomic confounding factors, such as chromatin context, 3D organisation, regulatory element redundancy/synergy, and poor scalability, such studies will be challenging and may require the development of new technologies.

## MATERIALS AND METHODS

### Selection of cCREs and promoters

For the design of the libraries we selected the cCREs and promoters from three TADs centered around each of the *Klf2, Nanog* and *Tfcp2l1* genes, using TAD coordinates from [54]. cCREs were selected based on DNAse hypersensitivity mapping data from mESCs in both 2i+LIF [38] and serum [5] culturing conditions, which we reprocessed and aligned to the mm10 genome build. DNAse hypersensitivity sites (DHSs) were called using Homer v4.10 with default parameters and peak style “factor”. We defined cCREs as 450 bp windows centered on each peak. For promoters we used the Gencode mouse TSS annotation [35]. From each TSS we defined as promoters the −375 +75 bp region. If the promoter regions overlapped with any cCRE then the promoter was redefined as the 450 bp region surrounding the center of the intersection of both elements. PCR primers were designed for each cCRE and promoter using the batch version of Primer3 (BatchPrimer3 v1.0) [64] allowing for primers to be designed on the 50 bps of each end. This yielded PCR products of ~400 bp for each element.

### Upstream assay library generation

For each locus, cCREs and promoters were amplified from mouse genomic DNA (extracted fromE14TG2a mESCs, ATCC CRL-1821) by PCR using My-Taq Red mix (#BIO-25044; Bioline) in 384 well plates using automated liquid handling (Hamilton Microlab^®^ STAR). PCRs were checked on gel and had a success rate between 60 and 90% depending on the locus. Equal volumes (10ul) of the resulting PCR products were mixed, and the resulting pool was purified by phenol-chloroform extraction followed by gel purification (BIO-52059; Bioline). The purified DNA fragments were then blunted and phosphorylated using End-It DNA End-Repair Kit (#ER0720; Epicentre). Part of the repaired pool was set apart for cloning of singlet libraries. The remainder was self-ligated using Fast-link ligase (LK0750H; Lucigen), after which duplets of ~800bp were excised from agarose gel and purified (BIO-52059; Bioline). Singlet and duplet pools were A-tailed using using Klenow HC 3’→5’ exo- (#M0212L; NEB).

The SuRE barcoded vector was prepared as described [30]. Then singlet and duplet pools were separately ligated overnight into the SuRE barcoded vector using Takara ligation kit version 2.1 (#6022; Takara). Ligation products were purified using magnetic bead purification (#CPCR-0050; CleanNA). Next, 2 μl of the purified ligation products were electroporated into 20 μl of electrocompetent e. cloni 10G supreme (#60081-1; Lucigen). Each library was grown overnight in 500 ml of standard Luria Broth (LB) with 50 μl/ml of kanamycin and purified using a maxiprep kit (K210016, Invitrogen).

### Downstream assay library generation

The Downstream assay vector was based on a pSMART backbone (Addgene plasmid # 49157; a gift from James Thomson). It was constructed using standard molecular biology techniques and contains a green fluorescent protein (GFP) open reading frame followed by a barcode, and a psiCheck polyadenylation signal (PAS) introduced during barcodin, followed by the cloning site for inserts and a triple polyadenylation site (SV40+bGH+psiCheckPAS).

The 10 highest expressing promoters of the *Klf2* Upstream library were selected to be cloned into the Downstream assay vector at the promoter position. These Promoters were amplified by PCR and individually inserted by Gibson assembly (#E2611S; NEB) into the Downstream assay vector. Then each of the 10 constructs were transformed into standard DH5α competent bacteria (#C2987; NEB) grown overnight in in 500 ml of standard Luria Broth(LB) with 50 μl/ml of kanamycin and purified.

Each of these promoter-containing vectors was then barcoded similarly as the SuRE vector [30]. For this, we digested 10 μg of each vector with AvrII (#ER1561; Thermo Fischer) and XcmI (#R0533; NEB) and performed a gel-purification. Barcodes were generated by performing 10 PCR reactions of 100 μl each containing 5 μl of 10 μM primer 275JvA, 5 μl of 10 μM primer 465JvA and 1 μl of 0.1 μM template 274JvA (see **Supplementary Table 2** for oligonucleotide sequences). A total of 14 PCR cycles were performed using MyTaq Red Mix (#BIO-25043; Bioline), yielding ~30 μg barcodes. Barcodes were purified by phenol-chloroform extraction and isopropanol precipitation after which they were digested overnight with 80 units of NheI (#R0131S; NEB) and purified using magnetic bead purification (#CPCR-0050; CleanNA). Each vector variant and the barcodes were then ligated in one 100 μl reaction containing 3 μg digested vector and 2.7 μg digested barcodes, 20 units NheI (#R0131S; NEB), 20 units AvrII, 10 μl of 10× CutSmart buffer, 10 μl of 10 mM ATP, 10 units T4 DNA ligase (#10799009001 Roche). A cycle-ligation of six cycles was performed (10 min at 22 °C and 10 min at 37 °C), followed by 20 min heat-inactivation at 80 °C. The ligation reaction was purified by magnetic beads and digested with 40 units of XcmI (#R0533S; NEB) for 3 h, and size-selected by gel-purification, yielding ~1 μg barcoded vector for each variant.

### Inverse PCR and sequencing to link inserted elements to barcodes

We identified barcode–insert combinations in the plasmid libraries by inverse-PCR followed by sequencing as described [30]. In brief, the combination of barcode and element(s) was excised from the plasmid by digestion with I-ceuI; this fragment was circularised; remaining linear fragments were destroyed; and circular fragments were linearised again with I-sceI. These linear fragments were amplified by PCR with sequencing adaptors. The final product was sequenced on an Illumina MiSeq platform using 150 bp paired-end reads. This process was done separately for each of the libraries. In the singlet libraries the barcodes should be associated to only one insert and in the combinatorial libraries the barcodes should be associated with duplets.

### Linking barcodes to element singlets or duplets

For each library the iPCR data was locally aligned using bowtie (version 2.3.4) [65] with very sensitive parameters (--very-sensitive-local) on a custom bowtie genome. This custom genome was generated using bowtie. It consists of virtual chromosomes corresponding to each cCRE or a P from each locus. Bam alignment files were processed using a custom python script that identifies from read 1 the barcode and cCRE or P element, and from read 2 the cCRE or P element. In case of singlet libraries both reads should identify the same element, whereas in combinatorial libraries read 1 is derived from the barcode-proximal element and read 2 from the barcode distal element. In the combinatorial libraries we can not distinguish between a combination of one element with itself in the same orientation or a single element, therefore these were removed from combinatorial libraries. In the Downstream Assay both reads identify the only element cloned in the downstream position. If no element was found, the barcode was assigned as empty vector. The resulting barcode-to-element(s) lists were clustered using Starcode (version 1.1) [66] to remove errors from barcode sequencing. Finally, barcodes present in multiple libraries or matched with multiple element combinations were removed from the data.

### Cell culture and transfection

All experiments were conducted in E14TG2a mouse embryonic stem cells (mESC) (ATCC CRL-1821) cultured in 2i+LIFulturing media. 2i+LIF was made according to the 4DN nucleome protocol for culturing mESCs (https://data.4dnucleome.org/protocols/cb03c0c6-4ba6-4bbe-9210-c430ee4fdb2c/). The reagents used were Neurobasal medium (#21103-049, Gibco), DMEM-F12 medium (#11320-033, Gibco), BSA (#15260-037; Gibco), N27 (#17504-044; Gibco), B2 (#17502-048; Gibco), LIF(#ESG1107; Sigma-Aldrich), CHIR-99021 (#HY-10182; MedChemExpress) and PD0325901 (#HY-10254; MedChemExpress), monothioglycerol (#M6145-25ML; Sigma) and glutamine (#25030-081, Gibco). Monthly tests (#LT07-318; Lonza) confirmed that the cells were not contaminated by mycoplasma. Cells were transiently transfected using Amaxa nucleofector II, program A-30, and Mouse Embryonic Stem Cell Nucleofector™ Kit (#VPH-1001, Lonza). *Klf2* and *Nanog* loci Upstream assay libraries were mixed and transfected together, *Tfcp2l1* Upstream Assay libraries were transfected in separate experiments. All the Downstream assay sub-libraries were transfected as a mix. Three independent biological replicates were done for each library mix. For each biological replicate 16 million cells were transfected (4 million cells with 4 μg plasmid per cuvette)

### RNA extraction and cDNA sequencing

RNA was extracted and processed for sequencing as described [30] with a few modifications. Cells were harvested 24 h after transfection, resuspended in Trisure (#BIO-38032; Bioline) and frozen at −80 °C until further processing. From the Trisure suspension, the aqueous phase containing the RNA was extracted and loaded into RNA extraction columns (#K0732, Thermo Scientific). Total RNA was divided into 10 μl reactions containing 5 μg of RNA and was treated for 30 mins with 10 units of DNAse I (#04716728001; Roche). Then DNAse I was inactivated by addition of 1 μl of 25 mM EDTA and incubation at 70°C for 10 min.

For the Upstream Assay the cDNA was produced and amplified by PCR as described [30]. Per biological replicate 8 to 10 reactions were carried out in parallel in order to cover enough barcode complexity of the library. For the Downstream Assay the RNA was extracted and processed the same way until cDNA production Here, cDNA was produced using a specific primer (304JvA sequence in **Supplementary Table 2** for oligonucleotide sequences). Primer 304JvA introduces an adaptor sequence 5’ to the primer sequence which is targeted in the first PCR (see below) to ensure strand specific amplification of barcodes. Then cDNA was amplified in 2 steps (nested PCRs) in order to make the reaction strand-specific. The first PCR reaction was run for 10 cycles (1 min 96 °C, 10 times (15 s 96 °C, 15 s 60 °C, 15 s 72 °C)) using (index variants of) primers 285JvA (containing the S2, index and p7 adaptor) and 305JvA (targeting the adapter introduced by 304JvA). Each 20 μl RT reaction was amplified in a 100-μl PCR reaction with MyTaq Red mix. The second PCR reaction was performed using 10ul of the product of the previous reaction in 100 μl reactions (1 min 96 °C, 8×(15 s 96 °C, 15 s 60 °C, 15 s 72 °C)) using the same index variant primer and primer 437JvA (containing the S1, and p5 adaptor). For both Upstream and Downstream assays, the resulting PCR products were sequenced on an Illumina 2500 HiSeq platform with 65bp single end reads.

### Plasmid DNA (pDNA) barcode sequencing

For normalisation purposes, barcodes in the plasmid pools were counted as follows. For both assays the process was the same. For each library 1 μg of plasmid was digested with I-sceI in order to linearise the plasmid. Then, barcodes were amplified by PCR from 50 ng of material using the same primers and reaction conditions as in the amplification of cDNA in the Upstream assay, but only 9 cycles of amplification were used (1 min 96 °C, 9 times (15 s 96 °C, 15 s 60 °C, 15 s 72 °C)). For each library, two technical replicates were carried out by using different index primers for each replicate. Samples were sequenced on an Illumina 2500 HiSeq platform with 65bp single end reads.

### Pre-Processing of cDNA and pDNA reads

For each replicate of each library pool transfection barcodes were extracted from the single end reads by using a custom python script that identifies the constant region after the barcode. Near-identical barcodes were pooled using Starcode (version 1.1) [66] to remove errors from barcode sequencing, and barcode counts were summarised. The process was the same for cDNA and pDNA counts and for Upstream and Downstream data.

### Post processing of cDNA and pDNA counts

For each transfection, barcodes identified in the cDNA were matched to the barcodes in the iPCR data, and all barcodes were counted in cDNA and pDNA replicates. Barcode counts were normalised to the total number of barcode reads from each sample. Activity per barcode was then calculated as a cDNA:pDNA ratio of normalised counts. Next, activities from multiple barcodes belonging to the same element singlet or combination were averaged, requiring a minimum of 5 barcodes per singlet or combination and at least 8 pDNA counts per barcode. The mean activity of each singlet or combination across replicates was calculated as the geometric mean of the three replicates.

### Calculation of boost indices

We initially calculated raw boost indices simply as a log_2_ ratio of the activity of each cCRE–P pair over the activity of the corresponding P alone. However, 20 negative controls that we included in the *Klf2* libraries, consisting of randomly generated DNA sequences of similar size and G/C content as the cCREs (**Supplementary dataset 1**), generally showed a negative boost index by this measure (median value −0.45 when inserted upstream) (**Figure S1D**). We therefore calculated corrected boost indices as the log_2_ ratio of cCRE-P activity over the median cCRE-P activity per promoter (**Figure S1D**). Importantly, in the *Klf2* library data this largely removed the negative bias that we observed with the negative controls; we thus assume that this correction is adequate and therefore also applied it to the boost indices obtained with the other libraries. For the analyses in **Figures 2–6** and **Supplementary figures 2–6** except **4A-B** the boost indices of cCREs were averaged over both orientations of the cCREs.

### Analysis of selectivity

We performed a Welch’s ANOVA (or Welch F-test) to assess the selectivity of each cCRE with more than 5 cCRE-P combinations. For this purpose, each replicate of each orientation of the cCRE-P was used as a datapoint and each cCRE-P combination was used as a group. P-values were corrected for multiple hypothesis testing using the Benjamini-Hochberg method and an FDR cutoff of 5% was chosen. The Welch F-test was chosen over the classic ANOVA due to heteroscedasticity of the data.

### TF motif Survey

We used a custom TF motif database provided by the lab of Gioacchino Natoli containing 2,448 TF motifs which was built on top of a previously published version [67] (Dataset composition and sources available at ##GitHub-url). TF motifs were filtered for expression of TFs in mESCs cultured in 2i+LIF according to published RNA-seq (higher expression than 1RPM) [38]. We scored presence or absence of a TF motif in each cCRE using FIMO (MEME suite, version 5.0.2). We then searched for motifs associated with (1) general enhancer activity, (2) self-compatibility and (3) duplets of self-compatible motifs. In (1), for each TF motif we compared the general cCRE-P population to combinations where the TF motif was present at the cCRE. In (2), for each TF motif we compared the cCRE-P combinations where the TF motif was present at the cCRE to the combinations where it was present at both the cCRE and the promoter. In (3), we took all the significant TF motifs at a 1% FDR and an effect size higher than 0.1 (n=66). Then we tested all pairwise non-repeated TF motif duplets. Per TF motif duplet we compared the cCRE-promoters where both TF motif were present at the cCRE to the combinations where both were present at both the cCRE and the promoter. In all comparisons a Wilcoxon test was applied to the boost indices of each group and the effect size was calculated a difference of median boost indices. In each analysis p-values were corrected for multiple hypothesis testing using the Benjamini-Hochberg method. We required a minimum of 50 combinations per group.

### Micro-C data correlation

Micro-C data was obtained from [55]. Contact scores between cCRE-P pairs were averaged across bins overlapping a +-500 bp window from the location of each element using 400 bp bins.

### Data analysis and data availability

All data analysis was performed in R [68]. Code of data processing pipelines and analysis scripts are available at ##Github-url. Raw and processed data are available at GEO (accession nr GSE186265). Processed datasets and pipeline output files are available at OSF (##OSF-ur).

## Author contributions

M.M.A, F.C. and B.v.S. designed the study. M.M.A and F.C. developed computational methods and performed analyses. M.M.A. and J.v.A. developed experimental methods. M.M.A. performed experiments. B.v.S. and M.M.A. wrote the manuscript, with input from F.C. and J.v.A. B.v.S. supervised the study.

## Acknowledgements

We thank Tao Chen for initial input on the study design, the NKI Genomics, Robotics and Research High Performing Computing facilities for technical support, Barak Cohen and his lab for insightful discussions, and Gioacchino Natoli and his lab for providing the TF motif database. Supported by ERC Advanced Grant 694466 (B.v.S.) and Swiss National Science Foundation postdoctoral fellowship P2EZP3_165206 (F.C). Oncode is partly funded by the Dutch Cancer Society KWF.

## Competing Interests

J.v.A. is founder of Gen-X B.V. and Annogen B.V. F.C. is a co-founder of enGene Statistics GmbH.

## SUPPLEMENTARY FIGURE LEGENDS

**Figure S1.**
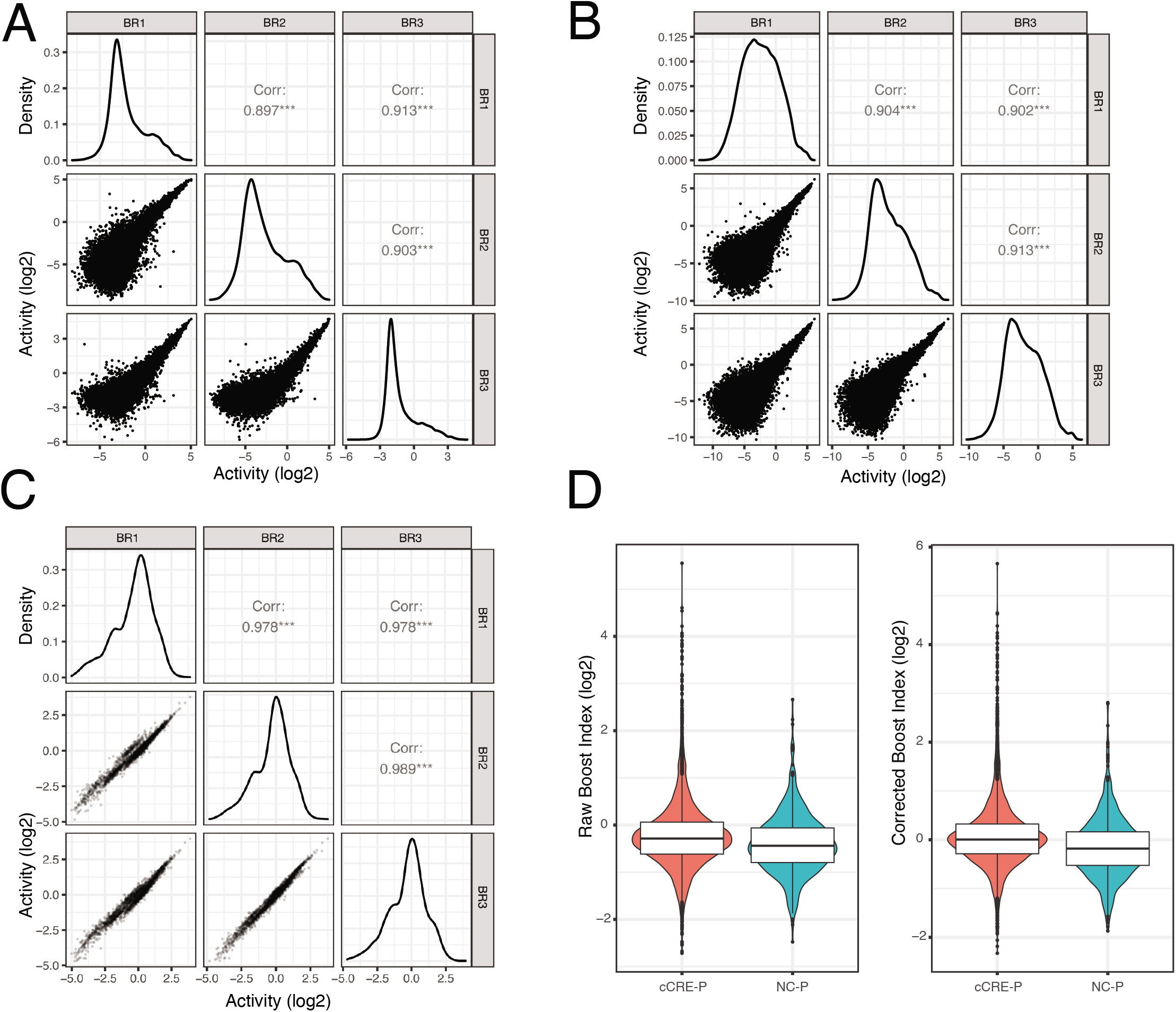
Reproducibility of data and boost index calculation. **(A-C)** Correlograms of the three biological replicates of each library pool. Lower left panels show pairwise scatterplots of the activities of all cCRE-P pairs per replicate. Middle panels show the density of data distribution in each replicate and upper right panels show the Pearson correlation coefficients. **A)** *Klf2* and *Nanog* Upstream libraries. **B)** *Tfcp2l1* Upstream library. **C)** *Klf2* Downstream libraries. **D)** Upstream assay boost index distributions for cCRE-P and negative controls – promoter (NC-P) combinations. Left panel: raw boost indices; right panel: boost indices after correction for negative bias (see Methods).

**Figure S2.**
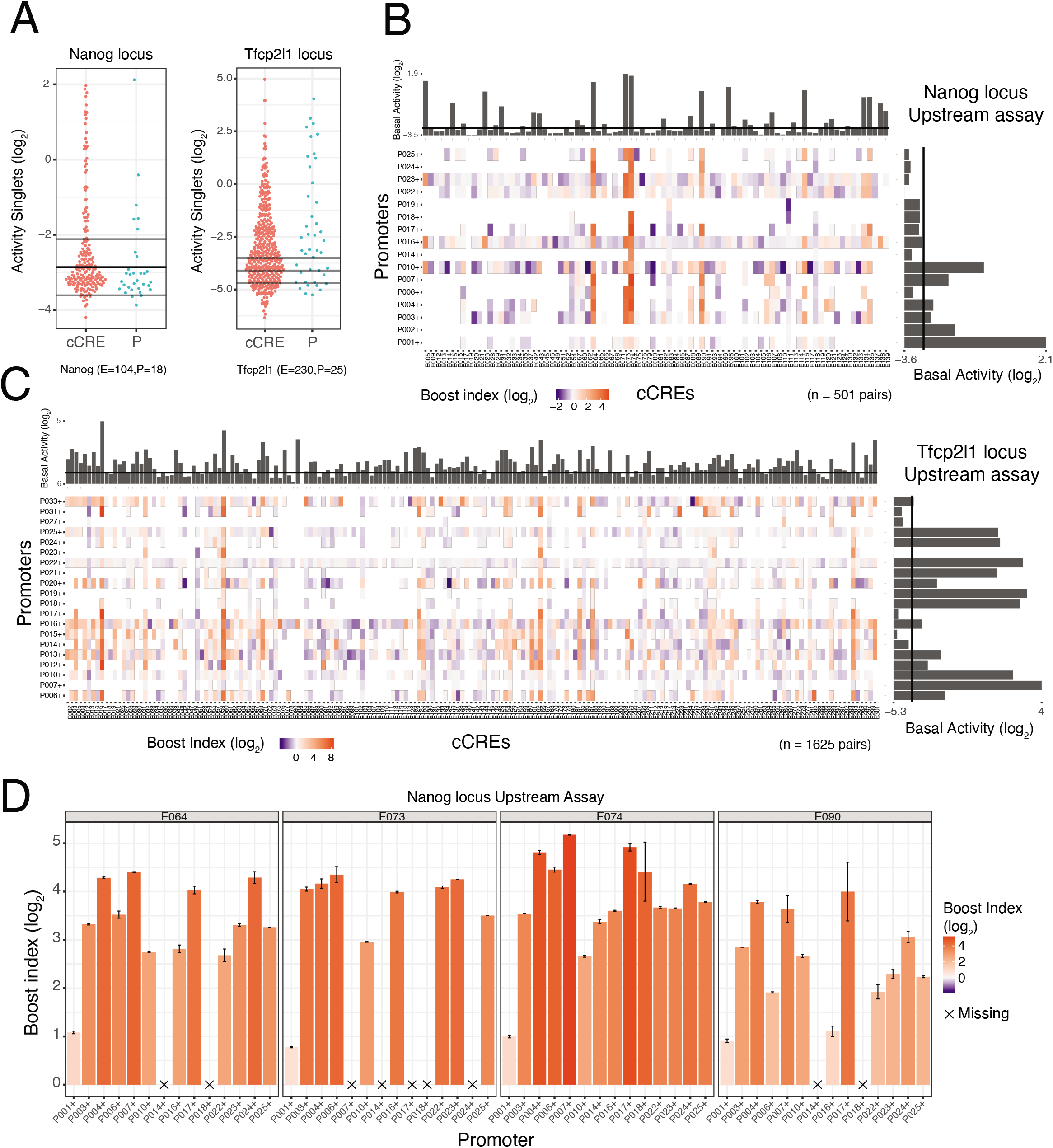
Element activities and boost indices obtained with *Nanog* and *Tfcp2l1* Upstream libraries. **A)** Transcriptional activities of cCREs and promoters. Each dot represents the mean activity of one singlet. Horizontal lines represent the average background activity of empty vectors (black line) plus or minus two standard deviations (grey lines). Elements with activities more than two standard deviations above the average background signal are defined as active. **B-C)** Boost index matrices for cCRE–P pairs from *Nanog* and *Tfcp2l1* loci (both Upstream assays). White tiles indicate missing data. Barplots on the right and top of each panel show basal activities of each tested P or cCRE, respectively, with the black line indicating the background activity of the empty vector. **D)** Examples of cCRE-P combinations for cCREs E064, E073, E074 and E090 of the *Nanog* locus. Barplots represent the mean boost index of each combination, vertical lines represent the standard deviation of each boost index. All data are averages over 3 independent biological replicates.

**Figure S3.**
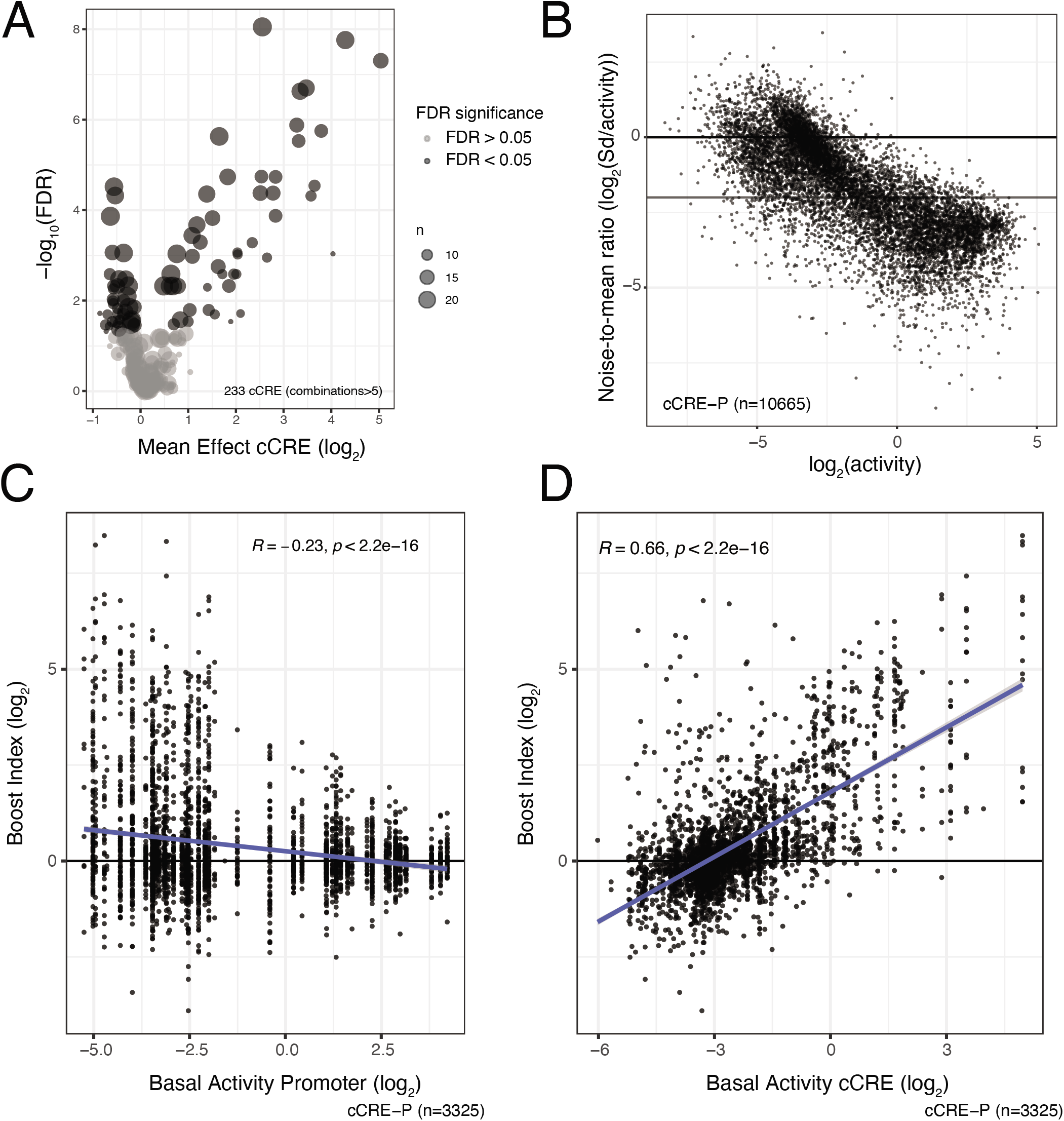
cCRE functional classification and activity influence on Boost indices. **A)** Volcano plot of cCREs associated with activation or repression across promoters. A Wilcoxon test is performed per cCRE comparing the boost indices of all the cCRE-P combinations of that cCRE against the rest of cCRE-P combinations. A minimum of 6 combinations is required per cCRE. P-values are corrected for multiple hypothesis testing using the Benjamini-Hochberg method (FDR). **B)** Relationship between noise-to-mean ratio (Standard Deviation/mean Activity) and mean activity of cCRE-Ps. Horizontal lines represent noise-to-mean ratios of 1 and of 4 in log2 scale. **C)** Relationship between boost indices and basal (singlet) P activity. Each column of dots shows the data of cCRE–P pairs for one P. Data are from Upstream assays of all three loci combined. **D)** Relationship between boost indices and basal (singlet) cCRE activity. All data are averages over 3 independent biological replicates.

**Figure S4.**
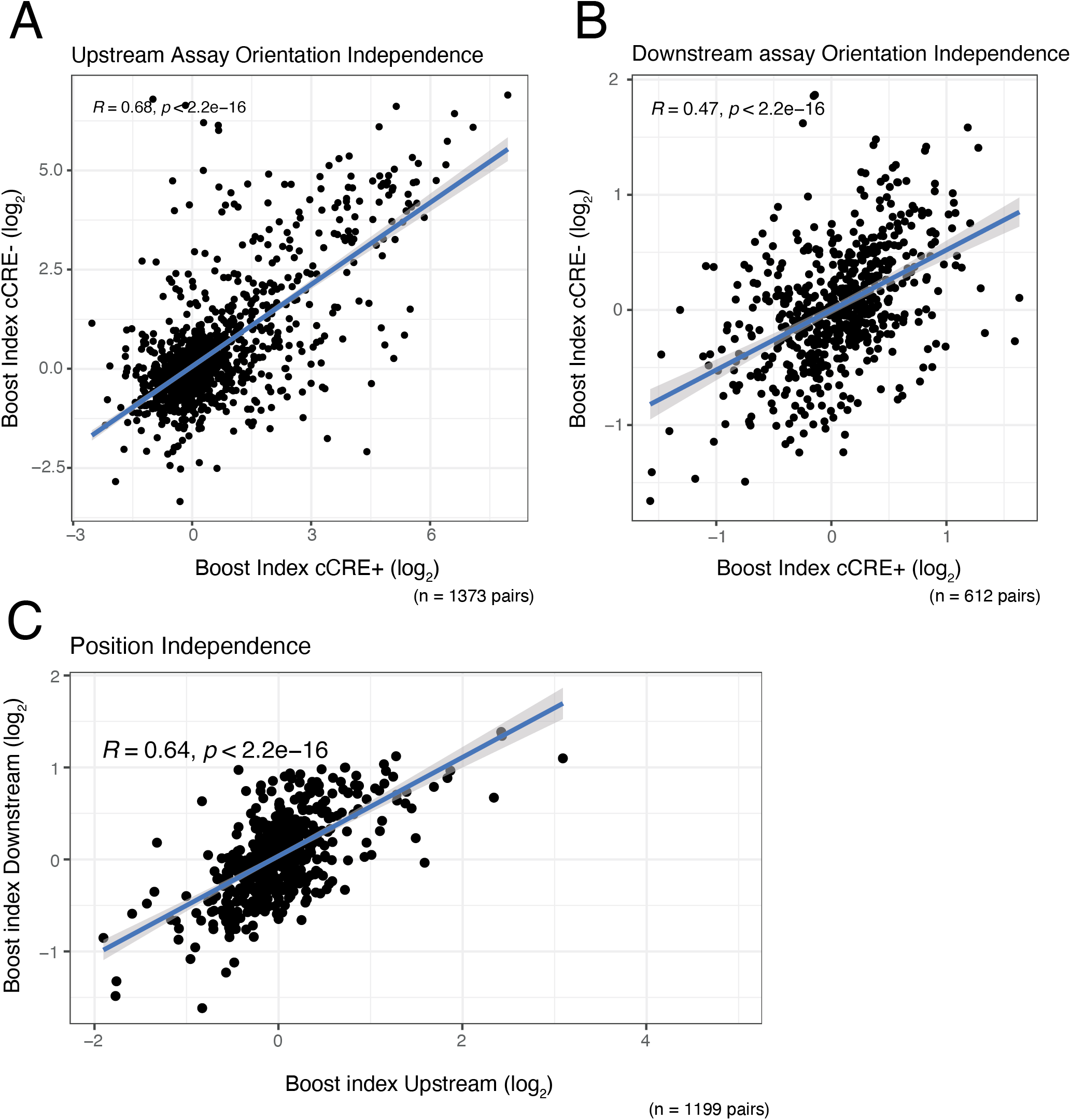
Orientation and position independence of cCREs. (**A-B)** Correlation between boost indices of both cCRE orientations of the same cCRE-P combination, in the **(A)** Upstream assay and **(B)** Downstream assay. Data are from the *Klf2* locus libraries. Note that “+” and orientations are arbitrary labels, because cCREs do not have an intrinsic orientation. **(C)** Correlation between boost indices of cCRE-P combinations shared between the Upstream and Downstream assays of the *Klf2* locus. In all panels R is the Pearson correlation coefficient. All data are averages over 3 independent biological replicates. In C Boost indices are averaged over cCRE orientations.

**Figure S5.**
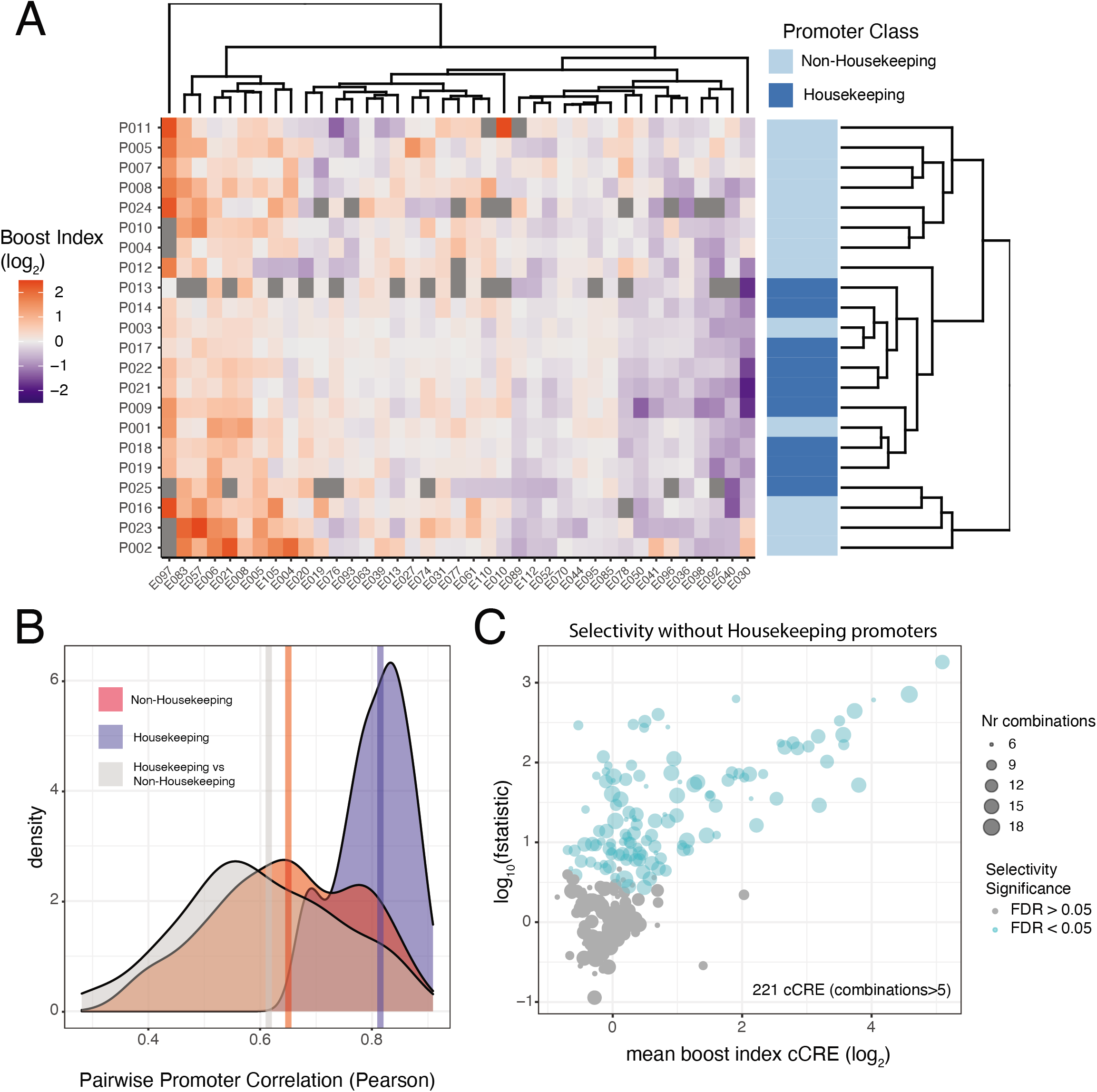
Housekeeping promoters show a distinct pattern of cCRE compatibility. **A)** Hierarchical clustering of the Upstream assay boosting matrix of the *Klf2* locus. In order to facilitate hierarchical clustering the matrix has been restricted to almost complete cases (cCREs >15 combinations) **B)** Density plot of pairwise Pearson correlation coefficients of the boost indices of *Klf2* locus promoters classified as either housekeeping or non-housekeeping [48]. Blue: correlations between all pairs of housekeeping promoters; red: all correlations between pairs of non-housekeeping promoters; grey: all correlations between one housekeeping and one non-housekeeping promoter. Vertical lines represent the median of each group. Unlike in (A), all promoters in the Upstream assay were included in this analysis. **C)** Results of selectivity analysis as performed in **Figure 4C**, but excluding housekeeping promoters. All data are averages over 3 independent biological replicates.

**Figure S6.**
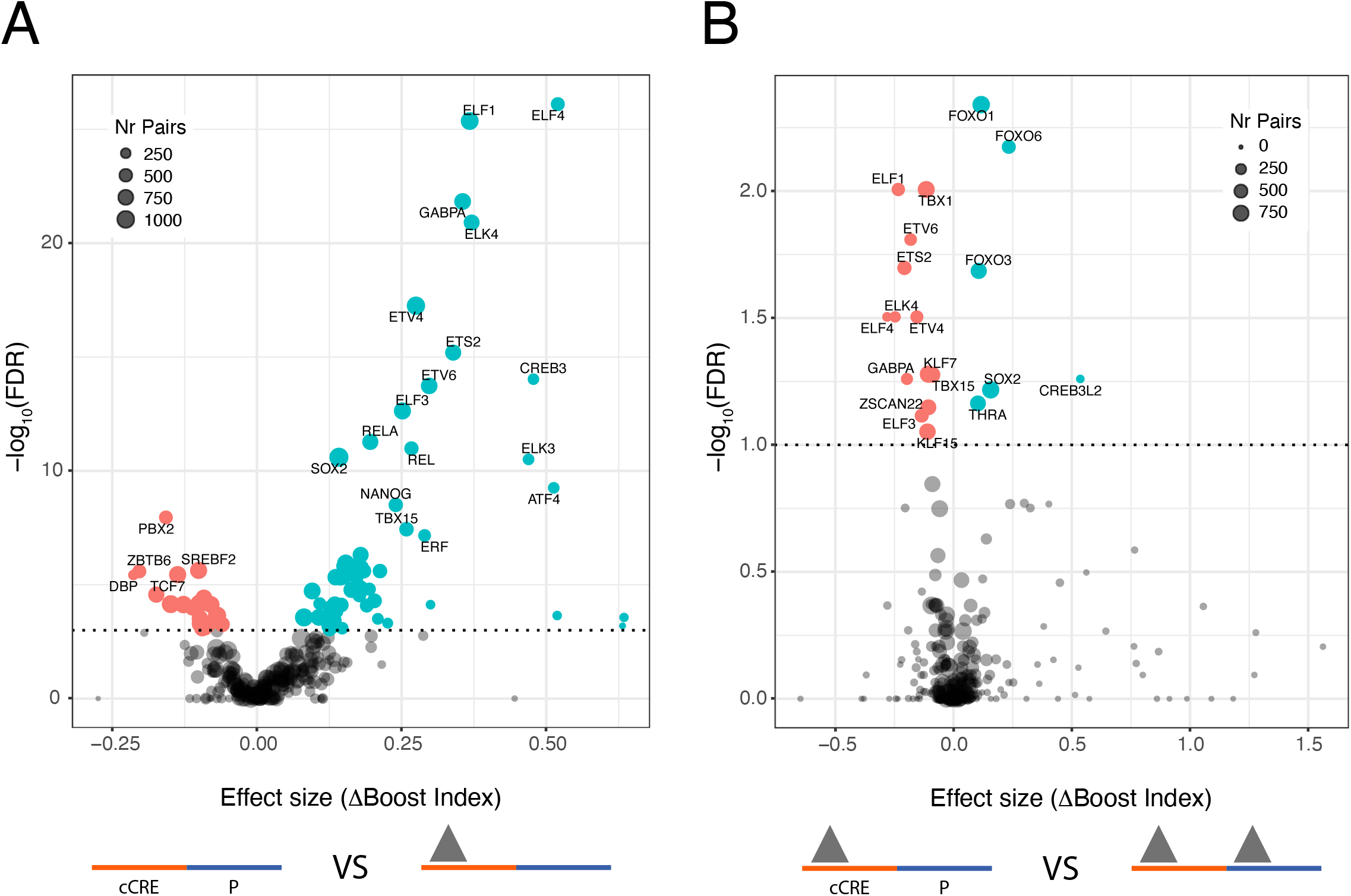
Identification of single TF motifs that correlate with boost indices. **(A)** TF motifs in cCREs associated (at 1% FDR cutoff) with activation (turquoise) or repression (red). **(B)** Motifs of putative self-compatible TFs, i.e. motifs that predict increased or reduced boosting indices when present both at the cCRE and P, compared to being present only at the cCRE. TF motifs associated with higher or lower boost indices at a 1% FDR cutoff are highlighted. We note that TF motifs with multiple hits from the same family, such as for ELK, FOXO and ELF factors, may in fact be due to the activity of one TF motif of that family [69].

## SUPPLEMENTARY TABLES

**Supplementary table 1.** Other combinations of cCRE and P elements in each MPRA library.

| Library | cCRE-cCRE | cCRE-cCRE (orientation-independent) | P-P | P-P (orientation-independent) | P-cCRE | P-cCRE (orientation-independent) |
| --- | --- | --- | --- | --- | --- | --- |
| Klf2 Upstream | 10626 | 4284 | 1335 | 441 | 4067 | 1439 |
| Nanog Upstream | 10536 | 4769 | 155 | 82 | 1511 | 713 |
| Tfcp2l1 Upstream | 44515 | 21149 | 626 | 274 | 5239 | 2386 |
| Klf2 Downstream | 0 | 0 | 420 | 225 | 0 | 0 |

**Supplementary table 2. Oligonucleotide and plasmid sequences**

**(supplementary file)**

## SUPPLEMENTARY DATASETS

**Data Set 1 Coordinates and sequences of cCREs and Promoters**

**Data Set 2 Activities of all cCRE-cCRE, cCRE-P, P-cCRE and P-P combinations Upstream assay**

**Data Set 3 Boost indices of cCRE-P combinations Upstream assay**

**Data Set 4 Boost indices of cCRE-P combinations Downstream assay**

## Notes

https://www.ncbi.nlm.nih.gov/geo/query/acc.cgi?acc=GSE186265

